# Intuitive interpretation of heterochromatin and euchromatin through rapid Hi-C analysis

**DOI:** 10.1101/2022.10.27.513973

**Authors:** Takashi Sumikama, Takeshi Fukuma

## Abstract

Hi-C is a technique that provides contact frequencies between pairs of loci on chromosomes. The conventional classification of heterochromatin and euchromatin based on Hi-C data is performed by principal component analysis; however, it requires long computational times and does not provide insight into the difference in contact frequencies between heterochromatin and euchromatin. Here, we propose a simple, intuitive and rapid method named the scaled contact number (SCN), which allows the contact frequencies to be visually interpreted and heterochromatin and euchromatin to be classified based on Hi-C results in a few minutes for long chromosomes at 1-kb resolution. The robustness of SCN was validated by confirming that SCN with reduced reads gives almost the same results as the original SCN. Overall, the approach described herein thus considerably decreases the time and computing power required to analyze Hi-C and further provides mechanistic insight indicating that euchromatin has more contacts than heterochromatin.

## Introduction

Hi-C technology has enhanced our understanding of chromosome structure and provided clues relating chromosome structure to functions^1^. A number of chromosome models that are compatible with Hi-C data have been suggested^1–7^. After normalization of the Hi-C matrix, chromosomes were classified into A (euchromatin) and B compartments (heterochromatin) according to the sign of the first or second eigenvector obtained by principal component analysis (PCA) of the Pearson correlation matrix. Normalization is critical to obtain correct eigenvectors^8^, and several different normalization methods to remove unwanted biases have therefore been proposed^9–14^. However, once it became possible to perform Hi-C at 1-kb resolution^15^, a technical problem in processing large amounts of Hi-C data emerged, as the diagonalization of the Pearson correlation matrix used in PCA requires very large amounts of memory and computational time^16^. Accordingly, fast and efficient methods of processing Hi-C data were developed^16–18^, but they still take a few days^19^. Another problem remains regarding the classification procedures themselves. Although the PCA-based classification of heterochromatin and euchromatin has worked well in practice, this mathematical procedure does not reveal the relation between the compartments and contact frequency. Moreover, the sign of the eigenvector after diagonalization is arbitrary^8^. Non-PCA-based classification methods have recently been suggested^19–21^, but the relation between the compartments and contact frequency remains unclear.

Here, to achieve rapid processing of a large amount of data without requiring a large amount of memory and to reveal the relation between the compartments and contact frequency, we propose a simple and intuitive classification method. The clarified difference in the interactions of compartments allows us to perform a rough classification by visually inspecting Hi-C results. This method requires only 3.4 MB of memory and takes less than 6 minutes to process one of the largest available Hi-C datasets (chr3 of the human B-lymphocyte cell line GM12878^15^) at 1-kb resolution using a standard laptop computer. Benchmarks of the processing time and comparison with CscoreTool^19^ are shown in Supplementary Fig. 1.

## Results

To test our classification method against the conventional method^1^, we first performed PCA on GM12878 (restriction enzyme: MboI)^15^, for which Hi-C was obtained at several resolutions, including 1 kb. The Hi-C matrix of chr14 at a 100-kb resolution and the corresponding eigenvectors obtained via the conventional procedure^1^ are shown in Fig. 1a. The scaled count number (SCN), a metric summing each column of Hi-C matrices developed here (see Method), is plotted at the bottom of Panel (a). SCN is very similar to the eigenvector, suggesting the accuracy of the new classification. The similarity is quantified later. This procedure allows us to calculate SCN at the highest resolution (1 kb) without requiring high computational power or large amounts of memory. SCNs of chr14 at 1, 5, 10, 25, 50 and 100 kb are shown (Supplementary Fig. 2). There is large fluctuation in the SCNs at 1 and 5 kb, and it is not clear whether this fluctuation is noise or reflects real local differences between heterochromatin and euchromatin. The fluctuation is suppressed at 10 kb, and the SCNs at 25 and 50 kb are essentially the same as those at 100 kb.

**FIGURE 1.**
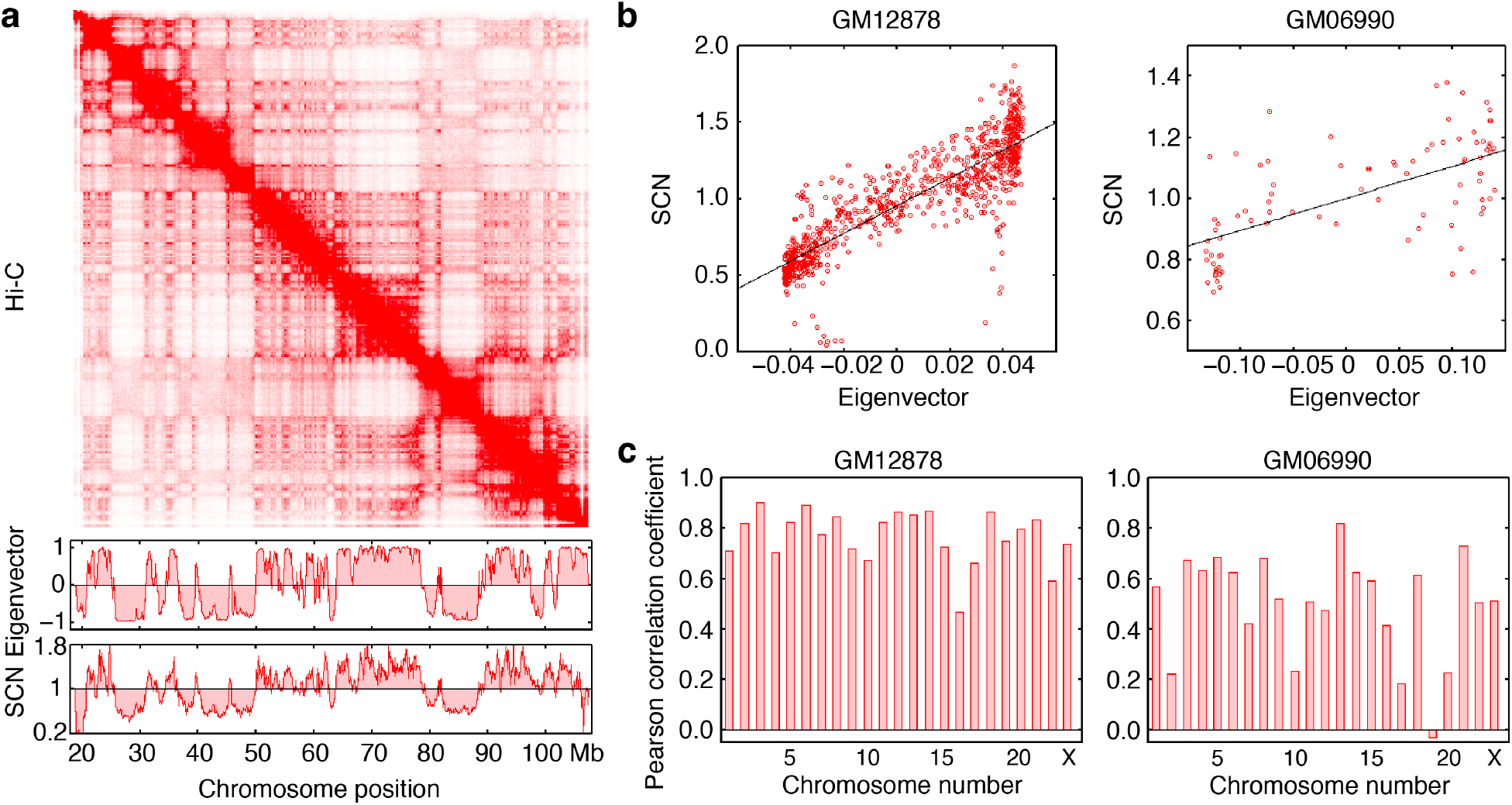
Comparison of eigenvectors and scaled count numbers (SCNs). (a) Hi-C matrix of chr14 of GM12878 B-lymphocyte cells at a 100-kb resolution, with eigenvectors and SCNs plotted below. (b) Plots of SCNs against eigenvectors. Data on GM12878 (chr14 at 100-kb resolution, restriction enzyme: MboI) and GM06990 (chr14 at 1-Mb resolution, restriction enzyme: HindIII) B-lymphocyte cells are also shown. Black lines are linear fits to the data. (c) Pearson correlation coefficients between SCNs and eigenvectors for all chromosomes in GM12878 and GM06990 B-lymphocyte cells.

To quantify the correlation between SCNs and eigenvectors, their relationship was plotted, and the results showed a good correlation (Fig. 1b). This was also the case for the low-resolution (1 Mb) Hi-C data obtained from another human B-lymphocyte cell line, GM06990 (restriction enzyme: HindIII)^1^ (Supplementary Fig. 3 for all the chromosome data), and higher resolution data (Supplementary Figs. 4a and 4b). The Pearson correlation coefficients between the SCNs and eigenvectors were high at 5-100 kb (Fig. 1c left and Supplementary Fig. 4c) and moderate at 1 Mb (Fig. 1c right), indicating the accuracy of SCN analysis.

We also compared the neighboring region contact index (NCI), a metric recently suggested by Fujishiro and Sasai for evaluating the correlation of contacts of a locus with its neighbours^20^, with the eigenvectors. The NCIs showed a high correlation with the eigenvectors at 100 kb but almost no correlation at 1 Mb (Supplementary Fig. 5). Therefore, we concluded that SCN is a more robust and meaningful measure than NCI.

To reveal the biological meaning of SCN, the values of SCN when the eigenvectors were 0 were plotted. The SCN intercept was consistently found to be 1 (Fig. 2a and Supplementary Fig. 4d), independent of the chromosome number or resolution. Thus, chromosome regions with an SCN > 1 or < 1 correspond to euchromatin and heterochromatin, respectively (Fig. 2b). Based on these results, heterochromatin and euchromatin can be roughly classified visually based on Hi-C results, without PCA, as illustrated in Fig. 2c. A higher than expected contact probability corresponds to euchromatin, which is open and more likely to contact other chromosome regions, while a lower probability corresponds to heterochromatin, which is packed and folded and thus contacts other chromosome regions infrequently (Fig. 2d). Thus, SCN is a clear indicator to distinguish between heterochromatin and euchromatin interactions.

**FIGURE 2.**
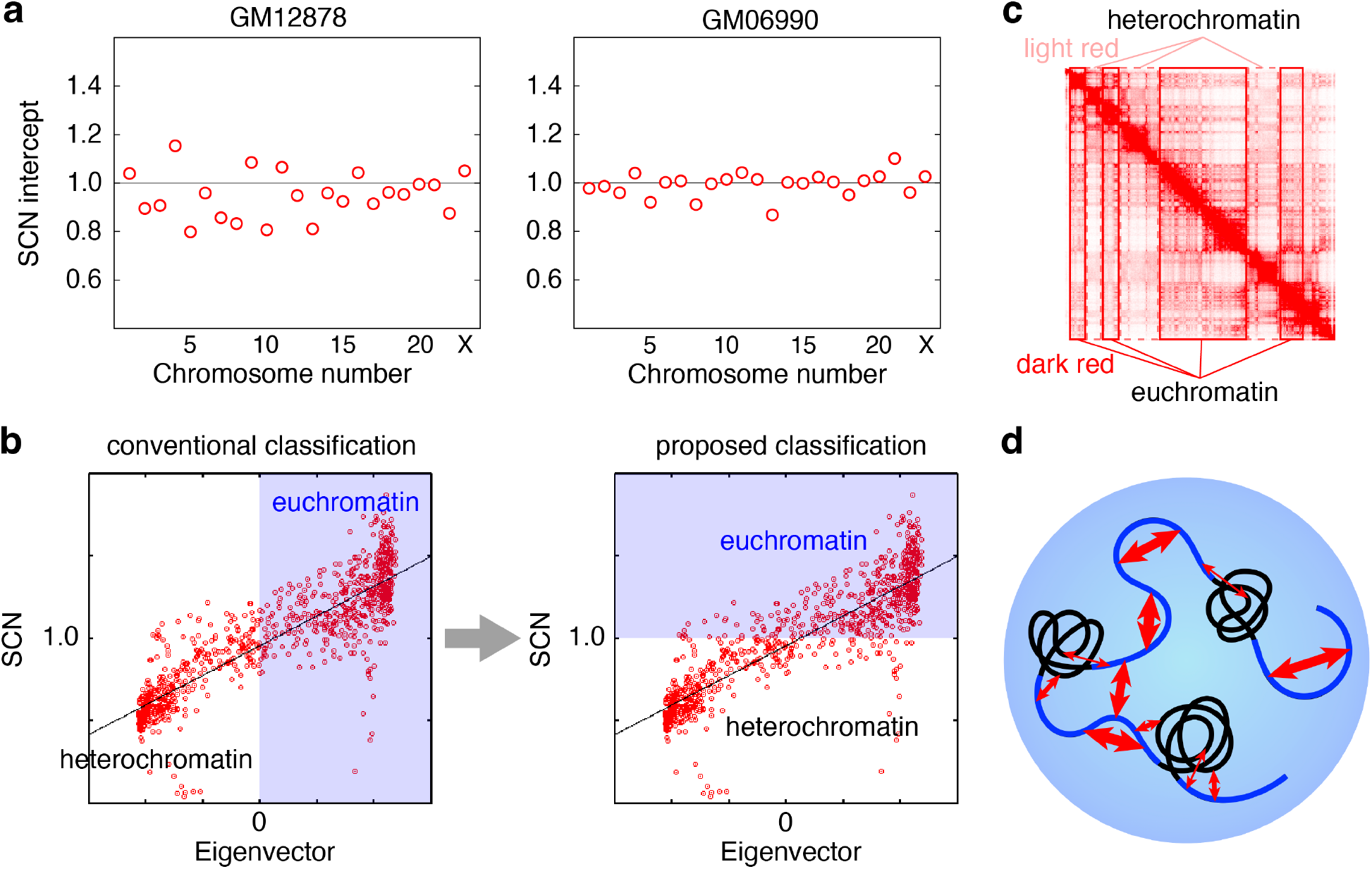
Biological meaning of SCN. (a) SCN intercepts (SCN values when the eigenvector is 0) of all chromosomes. (b) Conventional versus proposed classification of heterochromatin and euchromatin. (c) Visual classification of heterochromatin and euchromatin from Hi-C results. The dark red areas in the Hi-C matrix correspond to euchromatin, and the light red areas correspond to heterochromatin. (d) Schematic representation of heterochromatin (black) and euchromatin (blue) interaction. The thickness of the red arrows indicates the contact frequency inferred from SCNs.

It is important to further verify the robustness of this approach. As indicated earlier, the Pearson correlation coefficient for GM06990 is lower than that for GM12878, so one may assume that reads affect SCN (the reads for ch14 of GM06990 are 755k and those for GM12878 are 186 M). The reads of chr14 of GM06990 and GM12878 were randomly selected and reduced to 10%, 1%, and 0.1%. The SCNs calculated by using 10% and 1% data from GM12878 are indistinguishable from the original SCN, and the SCN of 0.1% data is very similar to the original SCN with some differences (Supplementary Fig. 6a). On the other hand, the SCN of 10% of the data from GM06990 is very similar to the original SCN with some differences, and the SCNs of 1% and 0.1% are different from the original SCN (Supplementary Fig. 6b). Accordingly, all SCNs with reduced data from GM12878 and SCNs with 10% data from GM06990 have similar Pearson correlation coefficients to the original SCN. Given that the lower number of reads (0.1% data of GM12878 (186k)) resulted in a higher Pearson correlation coefficient than the original GM06990 (755k), the correlation with the eigenvector does not depend only on the reads. Rather, SCN is influenced by the quality of Hi-C, since it is just a sum of the column of Hi-C matrices. In the Hi-C map of GM12878 (Fig. 1a), topologically associating domains (TADs) are clearly visible, while in the Hi-C map of GM06990 (see ref. 1), TADs are not as clear as in GM12878. Thus, SCNs are expected to have a high correlation with eigenvectors when applied to Hi-C data where TADs are clearly seen.

## Discussion

We developed a simple method for classifying heterochromatin and euchromatin based on Hi-C data that works even on laptops and can be readily implemented. Although it seems contrary to the conventional understanding, the results obtained via this method clearly demonstrated that the contact frequency of heterochromatin is lower than expected, while that of euchromatin is higher. This implies that euchromatin is flexible, easily interacts with other regions, and is accessible to the surface of heterochromatin. That is, euchromatin has frequent contacts in various locations. In contrast, heterochromatin is folded and less flexible than euchromatin and has a defined structure. Heterochromatin is in frequent or persistent contact within the defined structures, but chromosome parts inside the structures are difficult to access. Accordingly, heterochromatin has a lower contact frequency. Thus, SCN not only rapidly classifies heterochromatin and euchromatin but also offers insight into the different contact frequencies between heterochromatin and euchromatin.

There are several methods to identify TADs^22^. SCN appears to have some relation to TAD, as discussed above, and thus presumably contributes to improved identification methods. For example, a sudden change in SCN would be a sign of TAD boundaries. This classification method also has implications for developing simulation models. There were a few simulation models for heterochromatin and euchromatin^20,23,24^, where beads in the simulation model were assigned to heterochromatin or euchromatin using Hi-C data at 1-500 kb resolution. Using SCN, it is possible to assign beads at a resolution of 1-5 kb, corresponding to 5-25 nucleosomes, which is a requisite for future model development. At 1-kb or 5-kb resolution, large fluctuations were seen in SCN. Thus far, there is no information to determine if these fluctuations are noise or are truly capturing a heterochromatin/euchromatin difference, but such information is expected to be obtained by future simulation studies. Eventually, SCN will contribute to the development of a more realistic chromosome model implementing heterochromatin and euchromatin.

## Methods

Hi-C matrices (*m_ij_* record the counts of contacts between loci *i* and *j*. Accordingly, the sum of the column contents 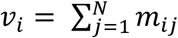, where *N* is the number of the row) gives the total number of contacts at locus *i*. To normalize these counts, the sums are scaled by the expected value 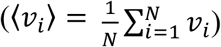. This is called the scaled contact number (SCN); that is, SCN = *v_i_*/〈*v_i_*〉. Columns with a sum of 0 counts were omitted from the analysis. A Fortran code to compute SCN reading by GM12878 is shown in the supplemental information.

## ACKNOWLEDGMENTS

T.S. thanks Dr. Hiratani and Dr. Miura at RIKEN for reading the manuscript. This work was supported by the World Premier International Research Center Initiative (WPI), MEXT, Japan; PRESTO (No. JPMJPR20K6), JST, Japan; and JSPS KAKENHI (No. 21H05251 and No. 20H00345). The calculation was carried out on the supercomputers at the Research Center for Computational Science in Okazaki, Japan (Project: 22-IMS-C114).

## AUTHOR CONTRIBUTIONS

T.S. and T.F. conceived the work. T.S. developed the program code, performed calculations, and drafted the manuscript. T.F. supervised the project. All the authors reviewed and edited the manuscript.

## ADDITIONAL INFORMATION

**Corresponding Author**

* (TS), (TF)

## REFERENCES

(1) Lieberman-Aiden, E. et al. Comprehensive Mapping of Long-Range Interactions Reveals Folding Principles of the Human Genome. Science 326, 289–293 (2009).

(2) Zhang, B. & Wolynes, P. G. Topology, structures, and energy landscapes of human chromosomes. Proc. Natl. Acad. Sci. USA 112, 6062–6067 (2015).

(3) Di Pierro, M., Zhang, B., Lieberman Aiden, E., Wolynes, P., & Onuchic, J. N. Transferable model for chromosome architecture. Proc. Natl. Acad. Sci. USA 113, 12168–12173 (2016).

(4) Di Stefano, M., Paulsen, J., Lien, T. G., Hoving, E., & Micheletti, C. Hi-C-constrained physical models of human chromosomes recover functionally-related properties of genome organization. Sci. Rep. 6, 35985 (2016).

(5) Kumari, K., Duenweg, B., Padinhateeri, R. & Prakash, J. R. Computing 3D Chromatin Configurations from Contact Probability Maps by Inverse Brownian Dynamics. Biophys. J. 118, 2193–2208 (2020).

(6) Lin, X., Qi. Y., Lathan, A. P. & Zhang, B. Multiscale modeling of genome organization with maximum entropy optimization. J. Chem. Phys. 155, 010901 (2021).

(7) Oliveira Jr., A. B., Estrada, C. P., Lieberman Aiden, E., Contessoto, V. G., & Onuchic, J. N. Chromosome Modeling on Downsampled Hi-C Maps Enhances the Compartmentalization Signal. J. Phys. Chem. B 125, 8757–8767 (2021).

(8) Fortin, J.-P. & Hansen, K. D. Reconstructing A/B compartments as revealed by Hi-C using long-range correlations in epigenetic data. Genome Biol. 16, 180 (2015).

(9) Yaffe, E. & Tanay, A. Probalistic modeling of Hi-C contact maps eliminates systematic biases to characterize global chromosomal architecture. Nat. Genet. 43, 1059–1065 (2011).

(10) Imakaev, M. et al. Iterative correction of Hi-C data reveals hallmarks of chromosome organization. Nat. Methods 9, 999–1003 (2012).

(11) Knight, P. A. & Ruiz, D. A fast algorithm for matrix balancing. IMA J. Numer. Anal. 33, 1029–1047 (2013).

(12) Schmitt, A. D., Hu, M. & Ren, B. Genome-wide mapping and analysis of chromosome architecture. Nat. Rev. Mol. Cell Biol. 17, 743–755 (2016).

(13) Lyu, H., Liu, E. & Wu, Z. Comparison of normalization methods for Hi-C data. BioTechniques 68, 56–64 (2020).

(14) Zheng, Y. & Keleş, S. Nomalization and de-noising of single-cell Hi-C data with BandNorm and sdVI-3D. Genome Biol. 23, 222 (2022).

(15) Rao, S. S. P. et al. A 3D map of the human genome at kilobase resolution reveals principles of chromatin looping. Cell 159, 1665–1680 (2014).

(16) Servant, N. et al. HiC-Pro: an optimized and flexible pipeline for Hi-C data processing. Genome Biol. 16, 259 (2015).

(17) Heinz, S. Simple combinations of lineage-determining transcription factors prime *cis*-regulatory elements required for macrophage and B cell identities. Mol. Cell 38, 576–589 (2010).

(18) Zheng, Y. & Keleş, S. FreeHi-C simulates high-fidelity Hi-C data for benchmarking and data augmentation. Nat. Methods 17, 37–40 (2020).

(19) Zheng, X. & Zheng, Y. CscoreTool: fast Hi-C compartment analysis at high resolution. Bioinformatics 34, 1586–1570 (2018).

(20) Fujishiro, S. & Sasai, M. Generation of dynamic three-dimensional genome structure through phase separation of chromatin. Proc. Natl. Acad. Sci. USA 119, e2109838119 (2022).

(21) Chakraborty, A., Wang, J. & Ay, F. dcHiC: differential compartment analysis of Hi-C datasets. bioRxiv doi: 10.1101/2021.02.02.429297 (2022).

(22) Forcato, M. et al. Comparison of computational methods for Hi-C data analysis. Nat. Methods 14, 679–685 (2017).

(23) Falk, M. et al. Heterochromatin drives compartmentalization of inverted and conventional nuclei. Nature 570, 395–399 (2019).

(24) Ancona, M. & Brackley, C. A. Simulating the chromatin-mediated phase separation of model proteins with multiple domains. Biophys. J. 121, 2600–2612 (2022).

