## Supplemental Information for "Intuitive interpretation of heterochromatin and euchromatin through rapid Hi-C analysis"

[1]PRESTO, JST, Kawaguchi, Saitama 332-0012, Japan.

[2]Nano Life Science Institute (WPI-NanoLSI), Kanazawa University, Kanazawa 920-1192, Japan.

* (TS), (TF)

### A sample code in Fortran to compute SCN

```
program Hi_C2SCN
  implicit none
  character(256) :: filename
  integer(8) :: i_min,i_min2,i,j,di,i_max,N,Nomit,ierr
  real(8) :: summ,mij
  integer,allocatable,dimension(:) :: flag
  real(8),allocatable,dimension(:) :: v

  call getarg(1,filename)
  open (10,file=filename,action='read')

  i_min = 500000000 ; i_min2 = 600000000 ; i_max = 0
  do
    read (10,*,iostat=ierr) i,j
    if (ierr /= 0) exit

    if (i < i_min) then
      i_min2 = i_min ; i_min = i
    elseif (i < i_min2 .and. i /= i_min) then
      i_min2 = i
    endif

    if (i > i_max) i_max = i
    if (j > i_max) i_max = j
  enddo
  rewind (10)

  di = i_min2 - i_min
  write (0,*) 'resolution =',di / 1000 ,'kb'
  write (0,*) 'maximum =',i_max / 1000,'kb'

  N = i_max / di ! number of columns or rows
  write (0,*) '# of data =',N
  allocate (v(N)) ; v = 0.d0
  allocate (flag(N)) ; flag = 0

  summ = 0.d0 ! summation of matrix
  do
    read (10,*,iostat=ierr) i,j,mij
    if (ierr /= 0) exit
    if (i == 0 .or. j == 0) CYCLE

    i = i / di
    j = j / di

    if (i /= j) then
      v(i) = v(i) + mij
      v(j) = v(j) + mij
      summ = summ + 2.d0 * mij
    else
      v(i) = v(i) + mij
      summ = summ + mij
    endif
  enddo

  Nomit = 0 ! number of columns to omit from analysis
  do i=1,N
    if (v(i) /= 0.d0) CYCLE
    flag(i) = 1
    Nomit = Nomit + 1
  enddo
```

```
  write (6,'(a2,2i15,f15.2)') '# ',N,Nomit,summ
  do i=1,N
    if (flag(i) == 1) then
      write (6,*)
    else
      j = i * di
      write (6,'(i15,f15.10)') j,v(i)/summ*(N-Nomit)
    endif
  enddo

  stop
end program Hi_C2SCN
```

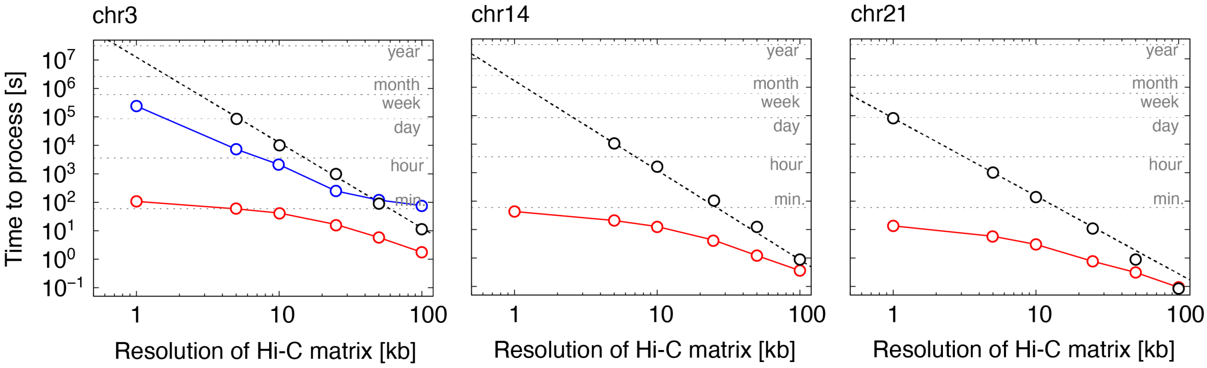


**Figure S1.** Processing speed benchmarks using an Intel Xenon Gold 2.4 GHz CPU. The black circles are the time required for principal component analysis written in Fortran, and the black dashed lines are fits. The red circles are the time required for SCN calculations. The processing time with CscoreTool[19] for chr1 of CM12878 using a 2.5 GHz CPU is shown in blue circles (data from ref. 19).

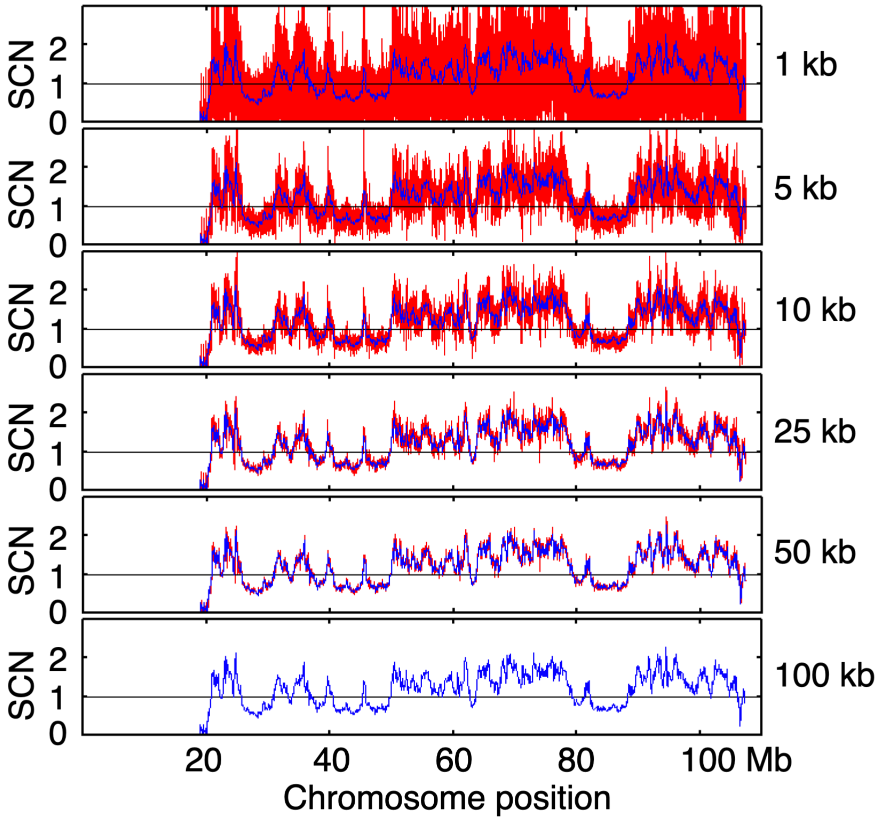


**Figure S2.** SCNs calculated for chr14 of GM12878 at a 1, 5, 10, 25, 50 and 100 kb resolution. For comparison, SCN at 100 kb is plotted in blue.

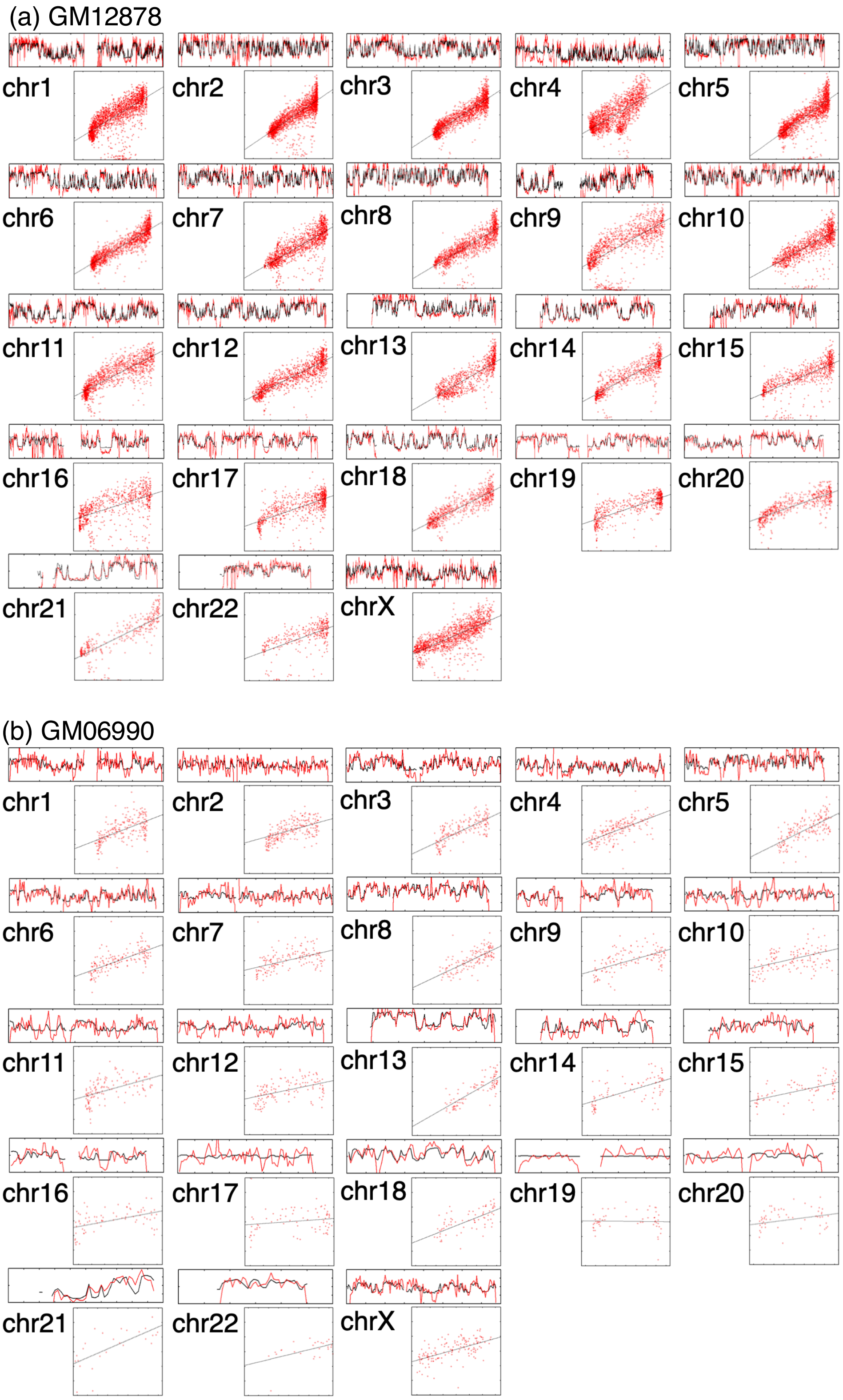


**Figure S3.** Comparison between SCNs and eigenvectors. (a) GM12878 at 100-kb resolution. (b) GM06990 at 1-Mb resolution. The upper panels show plots of SCNs (red) and eigenvectors (black) against chromosome loci. The ranges of the vertical axes indicating the SCNs of GM12878 and

GM06990 are 0.2-1.8 and 0.6-1.4, respectively. Lower panels are plots of SCNs against eigenvectors, with linear fits shown with black lines. The ranges of the vertical axes of the lower panels indicating the SCNs of GM12878 and GM06990 are 0-2 and 0.4-1.6, respectively. The ranges of the horizontal axes indicating the eigenvectors of the lower panels depend on the chromosome number.

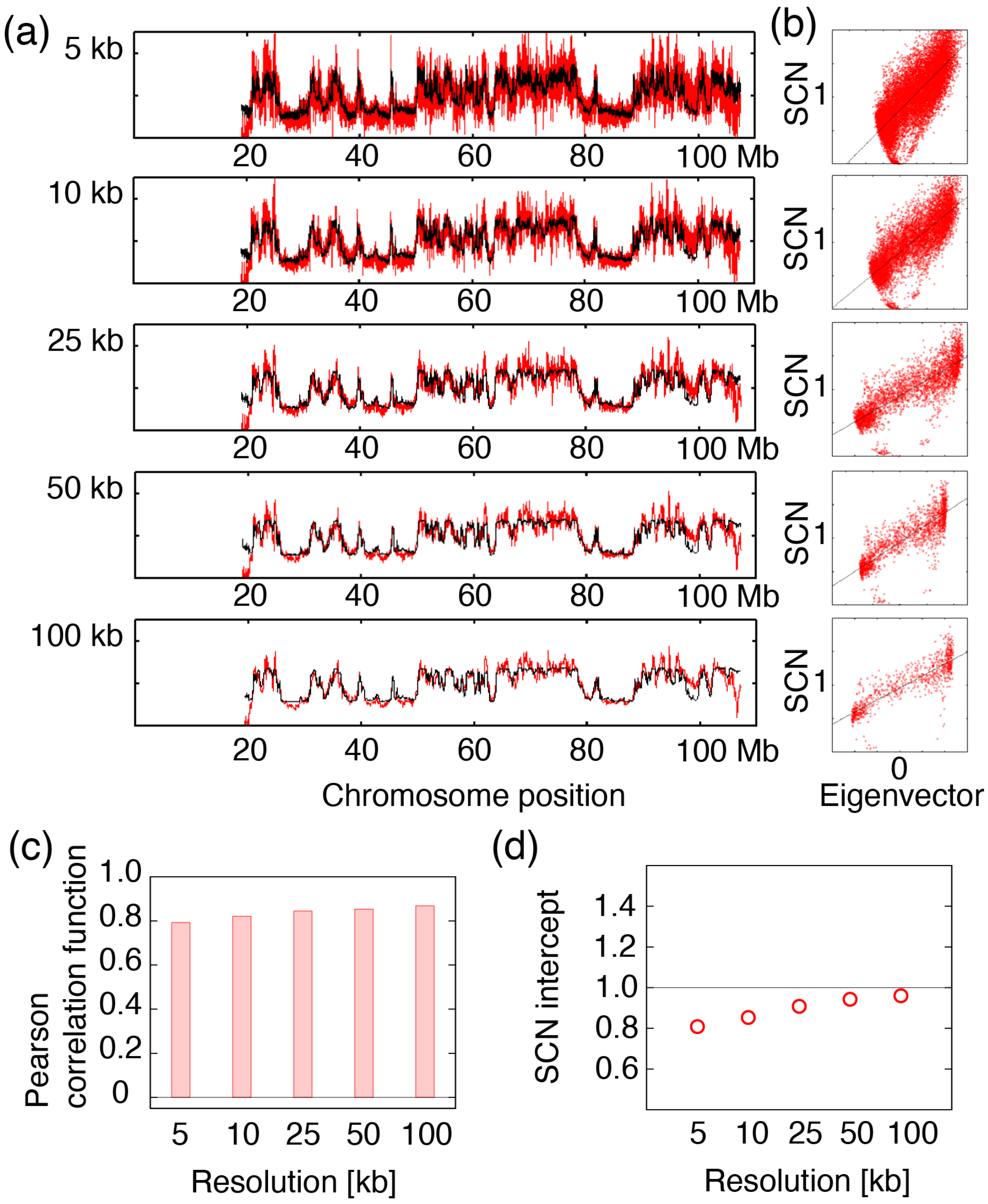


**Figure S4.** SCNs and eigenvectors for chr14 of GM12878 at a 5, 10, 25, 50 and 100 kb resolution. (a) Plots of SCNs (red) and eigenvectors (black) against chromosome loci. (b) SCNs against eigenvectors with linear fits shown by black lines. (c) Pearson correlation coefficients between SCNs and eigenvectors for chr14 at a 5, 10, 25, 50 and 100 kb resolution. (d) SCN intercepts (SCN values when the eigenvector is 0) of chr14 at a 5, 10, 25, 50 and 100 kb resolution.

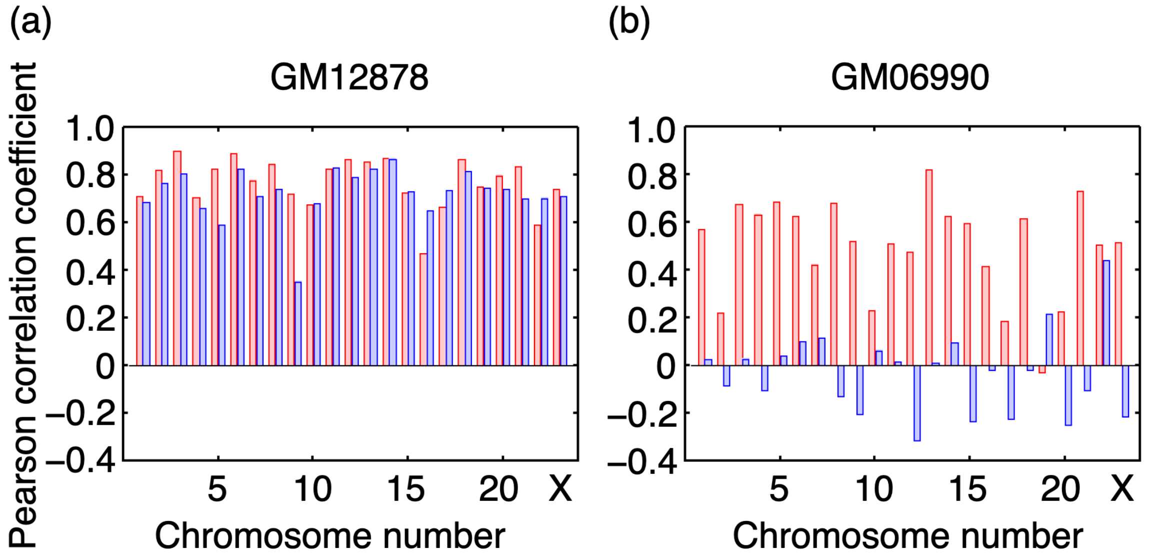


**Figure S5.** Pearson correlation coefficients between the SCNs and eigenvectors of all chromosomes (red) and those between NCIs and eigenvectors (blue). (a) GM12878 at 100-kb resolution. (b) GM06990 at 1-Mb resolution.

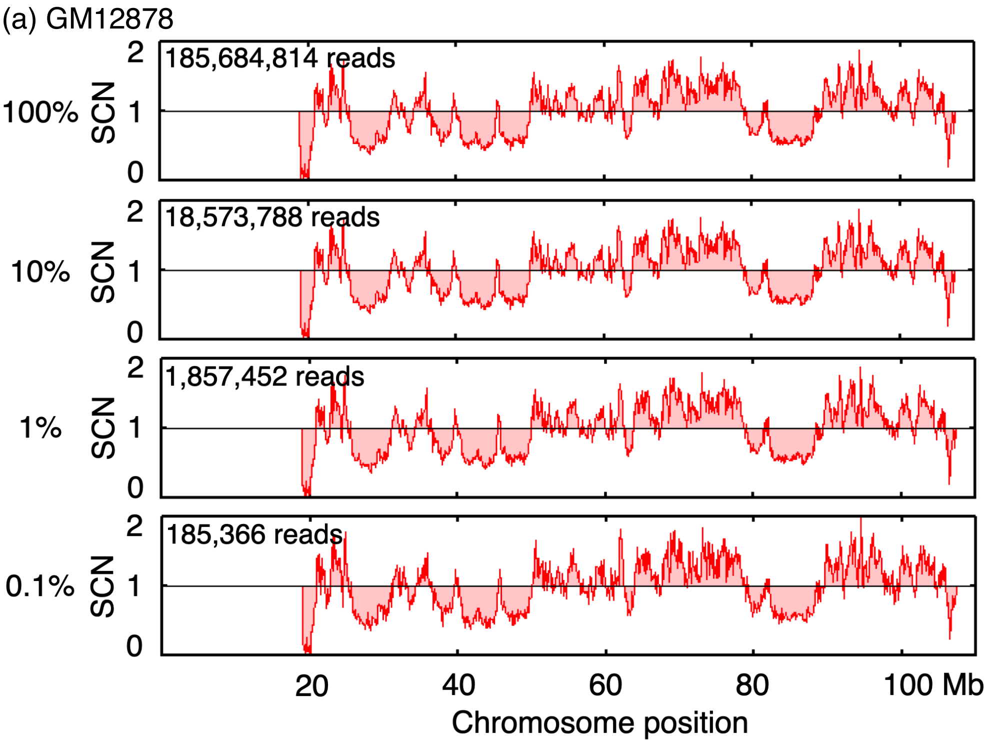


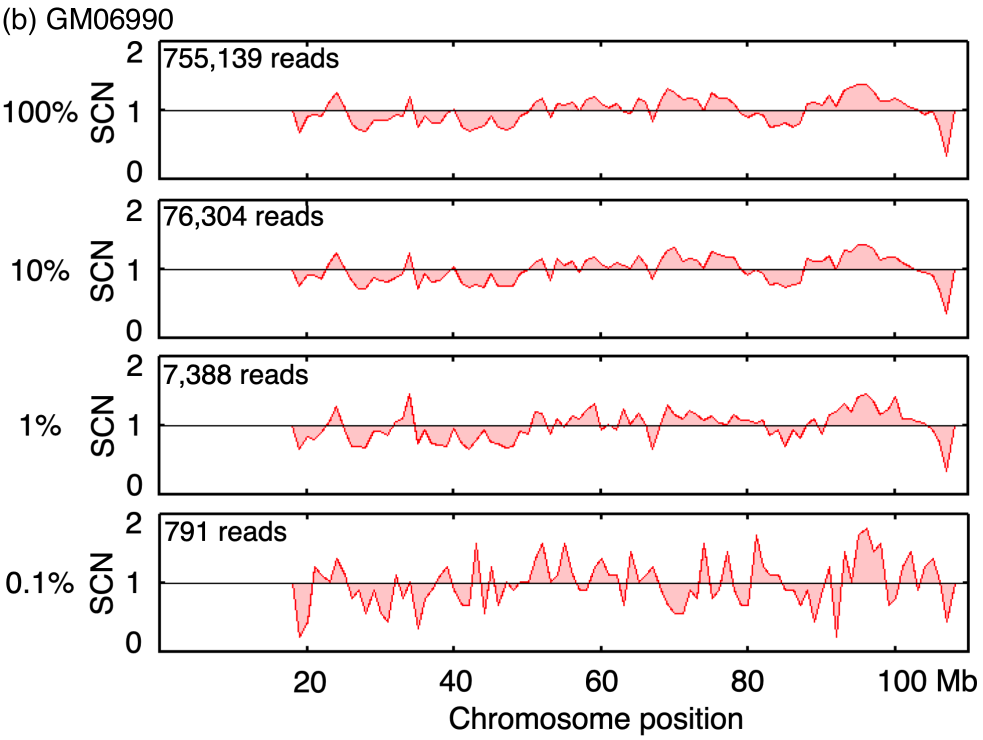


**Figure S6.** Comparison of SCNs when small fractions (10%, 1%, or 0.1%) of the reads were used. Each read is shown in each panel. (a) Chr14 of GM12878 at 100-kb resolution. (b) Chr14 of GM06990 at 1-Mb resolution.
